# Secondary bile acid ursodeoxycholic acid (UDCA) alters weight, the gut microbiota, and the bile acid pool in conventional mice

**DOI:** 10.1101/698795

**Authors:** Jenessa A. Winston, Alissa Rivera, Jingwei Cai, Andrew D. Patterson, Casey M. Theriot

**Affiliations:** Department of Population Health and Pathobiology, College of Veterinary Medicine, North Carolina State University, 1060 William Moore Drive, Raleigh, NC 27607; Department of Veterinary and Biomedical Sciences, The Pennsylvania State University, University Park, PA, 16802, USA

## Abstract

Ursodeoxycholic acid (commercially available as Ursodiol) is a naturally occurring bile acid that is used to treat a variety of hepatic and gastrointestinal diseases. Ursodiol can modulate bile acid pools, which have the potential to alter the gut microbiota community structure. In turn, the gut microbial community can modulate bile acid pools, thus highlighting the interconnectedness of the gut microbiota-bile acid-host axis. Despite these interactions, it remains unclear if and how exogenously administered ursodiol shapes the gut microbial community structure and bile acid pool. This study aims to characterize how ursodiol alters the gastrointestinal ecosystem in conventional mice. C57BL/6J wildtype mice were given one of three doses of ursodiol (50, 150, or 450 mg/kg/day) by oral gavage for 21 days. Alterations in the gut microbiota and bile acids were examined including stool, ileal, and cecal content. Bile acids were also measured in serum. Significant weight loss was seen in mice treated with the low and high dose of ursodiol. Alterations in the microbial community structure and bile acid pool were seen in ileal and cecal content compared to pretreatment, and longitudinally in feces following the 21-day ursodiol treatment. In both ileal and cecal content, members of the Lachnospiraceae family significantly contributed to the changes observed. This study is the first to provide a comprehensive view of how exogenously administered ursodiol shapes the gastrointestinal ecosystem. Further studies to investigate how these changes in turn modify the host physiologic response are important.

**Importance:** Ursodeoxycholic acid (commercially available as ursodiol) is used to treat a variety of hepatic and gastrointestinal diseases. Despite its widespread use, how ursodiol impacts the gut microbial community structure and bile acid pool remains unknown. This study is the first to provide a comprehensive view of how exogenously administered ursodiol shapes the gastrointestinal ecosystem. Ursodiol administration in conventional mice resulted in significant alterations in the gut microbial community structure and bile acid pool, indicating that ursodiol has direct impacts on the gut microbiota-bile acid-host axis which should be considered when this medication is administered.

**Bile Acid Abbreviations:** αMCA – α–Muricholic acid; βMCA –β–Muricholic acid; ωMCA –ω–Muricholic acid; CA – Cholic acid; CDCA – Chenodeoxycholic acid; DCA – Deoxycholic acid; GCDCA – Glycochenodeoxycholic acid; GDCA – Glycodeoxycholic acid; GLCA – Glycolithocholic acid; GUDCA – Glycoursodeoxycholic acid; HCA – Hyodeoxycholic acid; iDCA – Isodeoxycholic acid; iLCA – Isolithocholic acid; LCA – Lithocholic acid; TCA – Taurocholic acid; TCDCA – Taurochenodeoxycholic acid; TDCA – Taurodeoxycholic acid; THCA – Taurohyodeoxycholic acid; TUDCA – Tauroursodeoxycholic acid; TβMCA – Tauro-β-muricholic acid; TωMCA –Tauro ω-muricholic acid; UDCA – Ursodeoxycholic acid.

## Introduction

Bile acids are produced by host hepatocytes from cholesterol and are released into the gastrointestinal tract where they aid in the emulsification and absorption of dietary fat. Once host derived primary bile acids, namely cholic acid (CA) and chenodeoxycholic acid (CDCA) in humans, enter into the gastrointestinal tract the indigenous gut microbiota transforms them into secondary bile acids.^1,2^ Over 50 chemically distinct microbial derived secondary bile acids have been identified.^2^ Both primary and secondary bile acids can act as signaling molecules, exerting their effects by activating bile acid activated receptors, including G-protein coupled bile acid receptor 5 (TGR5) and the farnesoid X receptor (FXR)^3–5^. Examination of the gut microbiota-bile acid-host axis is growing in diverse fields including gastroenterology, endocrinology, oncology, immunology, and infectious disease.^1,3–13^

Ursodeoxycholic acid (UDCA) is a bile acid that has been medicinally utilized for over 2500 years.^14^ In humans, UDCA is considered a secondary bile acid derived from microbial conversion of the primary bile acid CDCA into lithocholic acid (LCA) and then into UDCA.^15^ However in other species, including mice, UDCA is a considered a host derived primary bile acid.^16–18^ The Food and Drug Administration (FDA) approved formulation of UDCA, or ursodiol, is used to treat a variety of diseases including: cholesterol gallstones, primary biliary cirrhosis, primary sclerosing cholangitis, nonalcoholic fatty liver disease, chronic viral hepatitis C, recurrent colonic adenomas, cholestasis of pregnancy, and recurrent pancreatitis.^6,19–27^ Ursodiol has vast beneficial effects (antichloestatic, antifibrotic, antiproliferative, and anti-inflammatory) but the major effect on bile acid physiology is an increase in hydrophilic bile acid pool by diluting the concentration of the hydrophobic toxic secondary bile acids, deoxycholic acid (DCA) and LCA.^6,28^

In healthy humans administered ursodiol (15 mg/kg/day) for 3 weeks, biliary and duodenal bile acid concentrations of UDCA and its conjugates (glycoursodeoxycholic acid, GUDCA and tauroursodeoxycholic acid, TUDCA) increased by 40% compared to baseline.^29^ A decrease in primary bile acids (CA and CDCA) and their glycine and taurine conjugates, as well as a decrease in the secondary bile acid DCA and its conjugates (glycodeoxycholic acid, GDCA and taurodeoxycholic acid, TDCA) was observed within biliary and duodenal bile.^29^ An increase in conjugates of the secondary bile acid LCA (glycolithocholic acid, GLCA and taurolithocholic acid, TLCA) were observed after UDCA treatment within biliary and duodenal bile samples.^29^ Ursodiol can alter liver and biliary bile acid pools, but gastrointestinal contents and feces have not been well studied, thus limiting our understanding of how ursodiol shapes the microbial niche and bile acid profiles within the gastrointestinal ecosystem.

Evidence is mounting that bile acids, through TGR5 and FXR signaling, are capable of altering the host physiologic response (recently reviewed in Wahlstrom et al.^3^ and Fiorucci et al.^4^). Bile acids can also directly and indirectly, through activation of the innate immune response, alter the gut microbial composition.^3,4^ Together, highlighting the interconnectedness and complexity of the gut microbiota-bile acid-host axis, and emphasizing the fact that exogenously administered bile acids will likely modulate this axis. Our rudimentary knowledge of how ursodiol modulates the gut microbial community structure, bile acid pool, and host physiology warrants further characterization to better understand the complex role of bile acids within the gastrointestinal ecosystem.

This study aims to define how ursodiol alters the gastrointestinal ecosystem in conventional mice. Mice were administered three different doses of ursodiol (50, 150, 450 mg/kg) via daily oral gavage for 21 days. The gut microbial community structure and bile acid pool were evaluated. Samples were obtained longitudinally in fecal samples and ileal and cecal content were collected pretreatment and after 21 days of ursodiol. Serum bile acid profiles were also evaluated after 21 days of ursodiol treatment. Collectively, ursodiol treatment resulted in biographically distinct alterations within the indigenous gut microbiota and bile acid metabolome in conventional mice. These findings support that ursodiol administration impacts the indigenous gastrointestinal ecosystem and thus modulates the gut microbiota-bile acid-host axis.

## Materials and Methods

### Ethical statement

The Institutional Animal Care and Use Committee (IACUC) at North Carolina State University College of Veterinary Medicine (NCSU) approved this study. The NCSU Animal Care and Use policy applies standards and guidelines set forth in the Animal Welfare Act and Health Research Extension Act of 1985. Laboratory animal facilities at NCSU adhere to guidelines set forth in the Guide for the Care and Use of Laboratory Animals. The animals’ health statuses were assessed daily, and moribund animals were humanely euthanized by CO_2_ asphyxiation followed by secondary measures (cervical dislocation). Trained animal technicians or a veterinarian performed animal husbandry in an AAALAC-accredited facility during this study.

### Animals and housing

C57BL/6J wildtype mice (females and males) were purchased from Jackson Laboratories (Bar Harbor, ME) and quarantined for 1 week prior to starting the Ursodiol administration to adapt to the new facilities and avoid stress-associated responses. Following quarantine, the mice were housed with autoclaved food, bedding, and water. Cage changes were performed weekly by laboratory staff in a laminar flow hood. Mice had a 12 hr cycle of light and darkness.

### Ursodiol dosing experiment and sample collection

Groups of 5 week old C57BL/6J WT mice (male and female) were treated with Ursodiol at three distinct doses (50, 150, and 450 mg/kg dissolved in corn oil; Ursodiol U.S.P., Spectrum Chemical, CAS 128-13-2) given daily via oral gavage for 21 days (Figure 1). Ursodiol dosing was adjusted once weekly, based on current weight. Two independent experiments were performed, with a total of n=8 mice (female/male) per treatment group. Mice were weighed daily over the course of the experiment. Fecal pellets were collected twice daily, flash-frozen and stored at −80°C until further analysis. A control group of mice were necropsied prior to initiating any treatments *(pretreatment* group). Necropsy was performed at day 21 in all Ursodiol treated mice. Gastrointestinal contents and tissue from the ileum and cecum were collected, flash frozen in liquid nitrogen, and stored at −80°C until further analysis. Serum and bile aspirated from the gallbladder was obtained flash frozen in liquid nitrogen, and stored at −80°C until further analysis.

**Figure 1:**
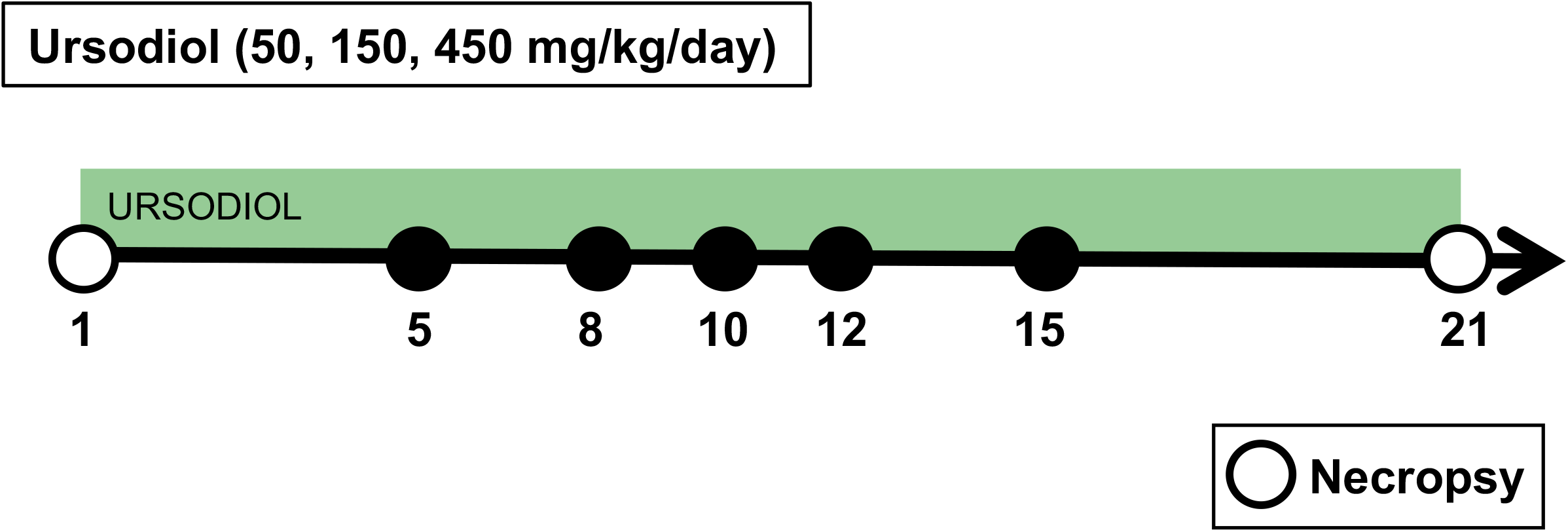
Mouse experimental design.Mouse experimental design.Mouse experimental design.Mouse experimental design. Groups of 5-week old C57BL/6J WT mice were treated with Ursodiol at three distinct doses (50, 150, and 450 mg/kg) given daily via oral gavage for 21 days. Fecal collection was performed twice daily throughout the experiment. Two independent experiments were performed, with a total of n = 8 (4 females/4males) mice per treatment group. Mice were monitored and weighed daily throughout the experiment. A control group of mice were necropsied prior to initiating any treatments (pretreatment group). Necropsy was performed at day 21 for all Ursodiol treated mice (open circles).

On several occasions, mice had evidence of corn oil within the oral cavity or on their muzzles immediately after the gavage. These mice were monitored closely for signs of aspiration pneumonia for 36 hr following this event. Two mice, one from the Ursodiol 50 mg/kg group and another from the Ursodiol 450 mg/kg group, inadvertently aspirated gavaged Ursodiol, containing corn oil, and subsequently developed respiratory distress within 12-24 hr following the aspiration event. The clinical signs were most consistent with lipid induced pneumonitis and both mice were humanely euthanized and excluded from the study.

### Targeted metabolomics of murine bile acid by UPLC-MS/MS

Targeted analysis of bile acids in ileal and cecal content, fecal pellets, serum, and bile were performed with an ACQUITY ultraperformance liquid-chromatography (UPLC) system using a C8 BEH column (2.1 × 100 mm, 1.7 μm) coupled with a Xevo TQ-S triplequadrupole mass spectrometer equipped with an electrospray ionization (ESI) source operating in negative ionization mode (All Waters, Milford, MA) as previously described.^30^ The sample was thawed on ice and 25 mg was added to 1 mL of precooled methanol containing 0.5 μM stable-isotope-labeled bile acids as internal standards (IS), followed by homogenization (1.0-mm-diameter zirconia/silica beads added) and centrifugation. Supernatant (200 μl) was transferred to an autosampler vial. 20 μL of serum was extracted by adding 200 μL pre-cooled methanol containing 0.5 μM IS. 5 μL of gall bladder bile was extracted with 500 μL pre-cooled methanol containing 0.5 μM IS. Following centrifugation, the supernatant of the extract was transferred to an autosampler vial for quantitation. Following centrifugation, the supernatant of the extract was transferred to an autosampler vial for quantitation. Bile acids were detected by either multiple reaction monitoring (MRM) (for conjugated bile acid) or selected ion monitoring (SIM) (for non-conjugated bile acid). MS methods were developed by infusing individual bile acid standards. Calibration curves were used to quantify the biological concentration of bile acids. Bile acid quantitation was performed in the laboratory of Dr. Andrew Patterson at Penn State University.

Random Forest analysis was performed in MetaboAnalyst 3.0 (http://www.metaboanalyst.ca/faces/ModuleView.xhtml).^31^ Briefly, the data were uploaded in the Statistical Analysis module with default settings and no further data filtering. Random Forest analysis Ward clustering algorithm and Euclidean distance were used to identify top bile acids within Ursodiol treatment groups. Heatmaps and box and whisker plots of bile acid concentrations, and nonmetric multidimensional scaling (NMDS) depicting the dissimilarity indices via Horn distances between bile acid profiles were generated using R packages (http://www.R-project.org).

### Illumina MiSeq sequencing of bacterial communities

Microbial DNA was extracted from murine fecal pellets and ileal and cecal tissue snips that also included luminal content using the PowerSoil-htp 96-well soil DNA isolation kit (Mo Bio Laboratories, Inc.). The V4 region of the 16S rRNA gene was amplified from each sample using a dual-indexing sequencing strategy.^32^ Each 20 μl PCR mixture contained 2 μl of 10× Accuprime PCR buffer II (Life Technologies), 0.15 μl of Accuprime high-fidelity Taq (catalog no. 12346094) high-fidelity DNA polymerase (Life Technologies), 2 μl of a 4.0 μM primer set, 1 μl DNA, and 11.85 μl sterile doubledistilled water (ddH2O) (free of DNA, RNase, and DNase contamination). The template DNA concentration was 1 to 10 ng/μl for a high bacterial DNA/host DNA ratio. PCR was performed under the following conditions: 2 min at 95°C, followed by 30 cycles of 95°C for 20 sec, 55°C for 15 sec, and 72°C for 5 min, followed by 72°C for 10 min. Each 20 μl PCR mixture contained 2 μl of 10× Accuprime PCR buffer II (Life Technologies), 0.15 μl of Accuprime high-fidelity Taq (catalog no. 12346094) high-fidelity DNA polymerase (Life Technologies), 2 μl of 4.0 μM primer set, 1 μl DNA, and 11.85 μl sterile ddH_2_O (free of DNA, RNase, and DNase contamination). The template DNA concentration was 1 to 10 ng/μl for a high bacterial DNA/host DNA ratio. PCR was performed under the following conditions: 2 min at 95°C, followed by 20 cycles of 95°C for 20 sec, 60°C for 15 sec, and 72°C for 5 min (with a 0.3°C increase of the 60°C annealing temperature each cycle), followed by 20 cycles of 95°C for 20 sec, 55°C for 15 sec, and 72°C for 5 min, followed by 72°C for 10 min. Libraries were normalized using a Life Technologies SequalPrep normalization plate kit (catalog no. A10510-01) following the manufacturer’s protocol. The concentration of the pooled samples was determined using the Kapa Biosystems library quantification kit for Illumina platforms (KapaBiosystems KK4854). The sizes of the amplicons in the library were determined using the Agilent Bioanalyzer high-sensitivity DNA analysis kit (catalog no. 5067-4626). The final library consisted of equal molar amounts from each of the plates, normalized to the pooled plate at the lowest concentration.

Sequencing was done on the Illumina MiSeq platform, using a MiSeq reagent kit V2 with 500 cycles (catalog no. MS-102-2003) according to the manufacturer’s instructions, with modifications.^32^ Libraries were prepared according to Illumina’s protocol for preparing libraries for sequencing on the MiSeq (part 15039740 Rev. D) for 2 or 4 nM libraries. The final load concentration was 4 pM (but it can be up to 8 pM) with a 10% PhiX spike to add diversity. Sequencing reagents were prepared according to Illumina’s protocol for 16S sequencing with the Illumina MiSeq personal sequencer.^32^ (Updated versions of this protocol can be found at http://www.mothur.org/wiki/MiSeq_SOP.) Custom read 1, read 2, and index primers were added to the reagent cartridge, and FASTQ files were generated for paired-end reads.

### Microbiome analysis

Analysis of the V4 region of the 16S rRNA gene was done using mothur (version 1.40.1).^32,33^ Briefly, the standard operating procedure (SOP) at http://www.mothur.org/wiki/MiSeq_SOP was followed to process the MiSeq data. The paired-end reads were assembled into contigs and then aligned to the SILVA 16S rRNA sequence database (release 132)^34,35^ and were classified to the mothur-adapted RDP training set v16^36^ using the Wang method and an 80% bootstrap minimum to the family taxonomic level. All samples with <500 sequences were removed. Chimeric sequences were removed using UCHIME.^37^ Sequences were clustered into operational taxonomic units (OTU) using a 3% species-level definition. The OTU data were then filtered to include only those OTU that made up 1% or more of the total sequences. The percentage of relative abundance of bacterial phyla and family members in each sample was calculated. A cutoff of 0.03 (97%) was used to define operational taxonomic units (OTU) and Yue and Clayton dissimilarity metric (θYC) was utilized to assess beta diversity. In addition to NMDS ordination, principle coordinate analysis (PCoA) biplots using Spearman correlation were used to examine difference in microbial community structures between Ursodiol treatments and compared to pretreatment. Standard packages in R (http://www.R-project.org) were used to create NMDS ordination on serial fecal samples.

### Statistical analysis

Statistical tests were performed using Prism version 7.0b for Mac OS X (GraphPad Software, La Jolla California USA) or using R packages (http://www.R-project.org). To assess weight loss a two-way ANOVA with Dunnett’s multiple comparisons post hoc test comparing Ursodiol treatment groups and untreated mice was performed. For microbiome analysis, analysis of molecular variance (AMOVA) was used to detect significant microbial community clustering of treatment groups in NMDS plots and principle coordinate analysis (PCoA) biplots using Spearman correlation were used to examine difference in microbial community structures between Ursodiol treatments and compared to pretreatment.^38^ For bile acid metabolome, a NMDS illustrates dissimilarity indices via Horn distances between bile acid profiles. To assess the comprehension bile acid profiles, a two-way ANOVA followed by Dunnett’s multiple comparisons post hoc test was used to compare Ursodiol treatment groups to pretreatment bile acid profiles. A Kruskal-Wallis one-way ANOVA test followed by Dunn’s multiple comparisons test was used to calculate the significant of individual bile acid within each Ursodiol treatment group compared to pretreatment. Statistical significance was set at a p value of < 0.05 for all analyses (*, p <0.05; **, p < 0.01; ***, p < 0.001; ****, p < 0.0001).

## Results

### Ursodiol treatment results in weight loss

C57BL/6J conventional mice were administered three different doses of ursodiol (50, 150, 450 mg/kg/day; denoted here on out as Ursodiol 50, Ursodiol 150, and Ursodiol 450 respectively) via oral gavage for 21 days (Figure 1). Mice were monitored and weighed daily. Mice in the 50 and 450 mg/kg ursodiol treatment groups sustained significant weight loss within a week of administration of Ursodiol compared to untreated mice (Figure 2A and 2C). For the Ursodiol 50 mg/kg treatment group, this weight loss persisted over the course of the experiment (Figure 2A). For the ursodiol 450 mg/kg treatment group, initially weight loss was noted during the first and third week of Ursodiol administration (Figure 2C). The ursodiol 150 mg/kg treatment group did not have significantly different weights compared to the untreated mice (Figure 2B). No other clinical signs were noted during Ursodiol administration. In general, mice tolerated daily gavage with diminishing stress related to the procedure over the course of the experiment.

**Figure 2:**
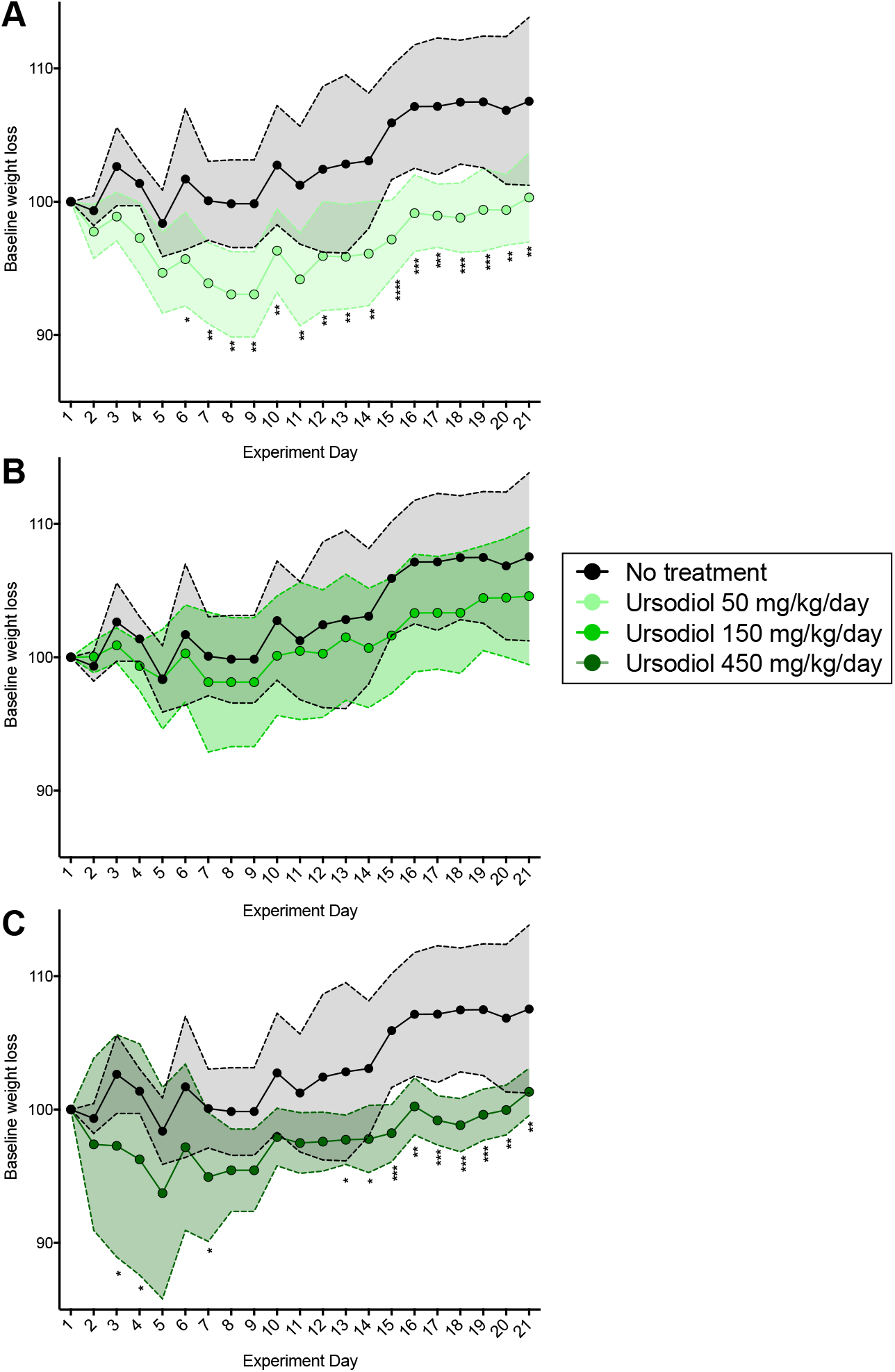
Weight loss observed with daily Ursodiol administration. **(A)** Weight loss in Ursodiol 50 mg/kg, **(B)** Ursodiol 150 mg/kg, and **(C)** Ursodiol 450 mg/kg treatment group compared to untreated mice. Statistical significance between Ursodiol treatment groups and untreated mice was determined by a two-way ANOVA with Dunnett’s multiple comparisons post hoc test. Shaded regions represent the standard deviations from the mean. For all graphs (*, p ≤ 0.05; **, p ≤ 0.01; ***, p ≤ 0.001; ****, p ≤ 0.0001). Data represents two independent experiments.

### Ursodiol alters the gut microbial community structure in conventional mice

Paired fecal samples were collected from the same mice serially over the 21-day experiment to facilitate simultaneous evaluation of the microbial community structure and bile acid metabolome. Mice were sacrificed at day 21 and gut content from the ileum and cecum were collected at necropsy, and stored for later analysis. 16S rRNA gene sequencing was performed to define the gut microbiota.

Within the ileum, the gut microbial community structure of the Ursodiol 150 and Ursodiol 450 treatment groups were significantly different from pretreatment (Figure 3A; AMOVA; p = 0.02 and p = 0.009, respectively). Bar plots were utilized to visualize relative composition of ileal microbial communities, which are different across each Ursodiol dose and compared to pretreatment (Figure 3C). However, the overall gut microbial community structure between treatments was not significantly different based on AMOVA. A biplot of the correlating OTUs towards PCoA axes 1 and 2 revealed OTU 109 (classified as Lachnospiraceae) as the only significant member contributing to ileal microbial community alterations seen with Ursodiol treatment (Figure S1A and Figure 3C).

**Figure 3:**
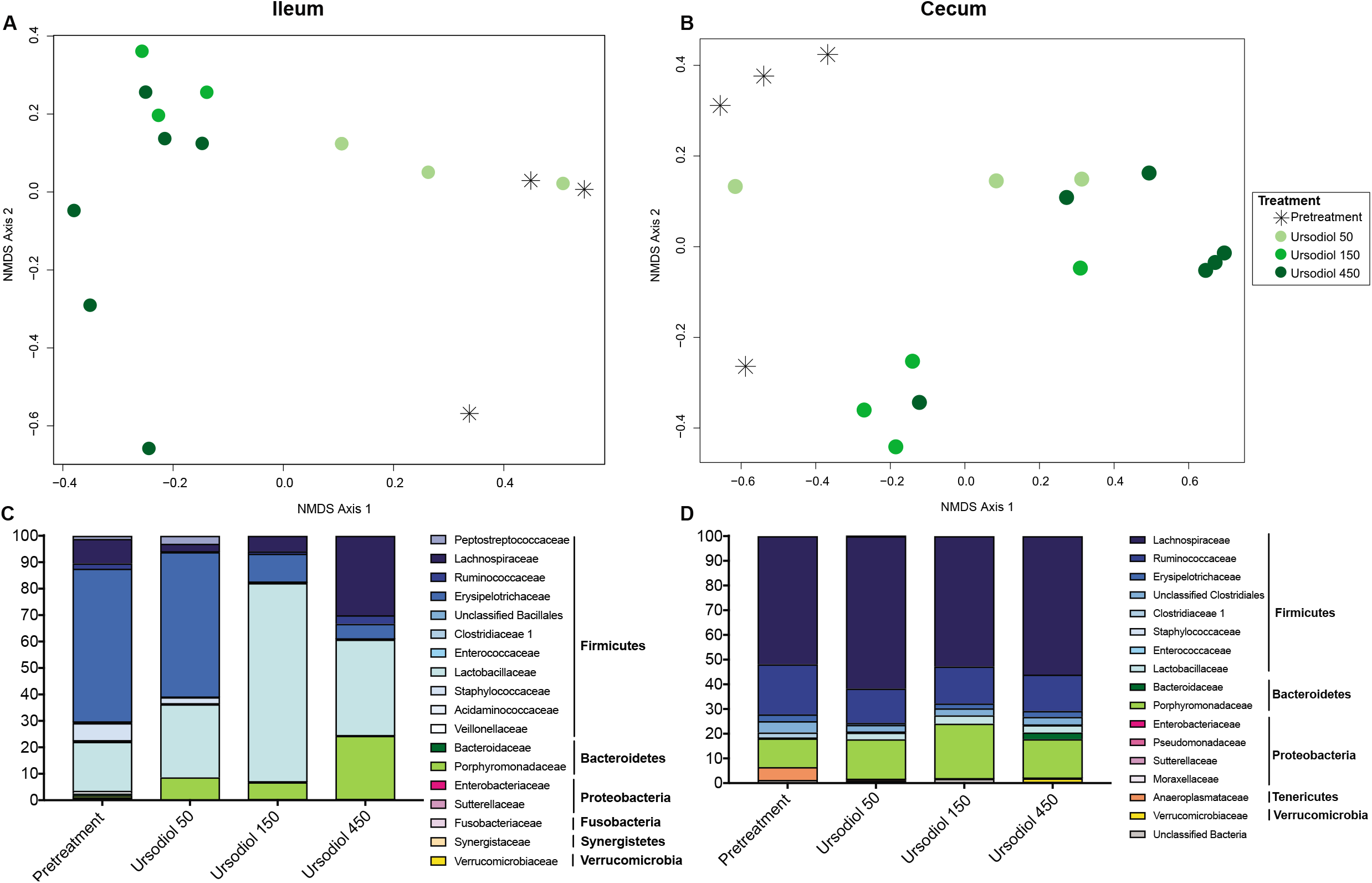
Alterations to the indigenous ileal and cecal microbiota associated with Ursodiol administration in conventional mice. NMDS ordination was calculated from Yue and Clayton dissimilarity metric (θ_YC_) on OTU at a 97% cutoff of **(A)** ileal and **(B)** cecal samples from pretreatment and Ursodiol treated mice. Statistical significance between Ursodiol treatment groups and pretreatment mice was determined by AMOVA. The composition of the **(C)** ileal and **(D)** cecal microbiota was visualized with bar plots of the family relative abundance for each treatment group (n=3 mice per treatment).

Within the cecum, the gut microbial community structure of the Ursodiol 450 treatment group was significantly different from pretreatment (Figure 3B; AMOVA; p = 0.002). Bar plots were utilized to visualize relative composition of cecal microbial communities, which were marginally different across each Ursodiol dose and compared to pretreatment (Figure 3D). In accordance, the overall gut microbial community structure between treatments was not significantly different based on AMOVA. A biplot of the top 10 OTUs towards PCoA axes 1 and 2 revealed OTU 86 (classified as Lachnospiraceae) as a significant member contributing to cecal microbial community alterations seen with Ursodiol treatment (Figure S1B).

Within the feces, the gut microbial community structures of all Ursodiol treatment groups were significantly different from pretreatment (Figure S1C; AMOVA; p = 0.004, p <0.001, p <0.001, respectively). A biplot of the top 10 correlating operational taxonomic units (OTUs) towards PCoA axes 1 and 2 revealed OTU 24 (classified as Ruminococcaceae) as a significant member contributing to fecal microbial community alterations seen with Ursodiol treatment over time and eight opposing OTUs (Figure S1C).

### Ursodiol alters the bile acid pool in conventional mice

To determine the extent that ursodiol alters the bile acid pool, assessment of 47 bile acids, was conducted on paired ileal, cecal, and fecal samples used in the preceding microbial community structure evaluation. In addition to NMDS ordination and comprehensive bile acid profile heatmaps, Random Forest analysis was applied to identify bile acids that are important for distinguishing between ursodiol treatments.

Ileal content bile acid profiles revealed segregation of the ursodiol 150 and ursodiol 450 treatments from pretreatment bile acid profiles (Figure 4A). A total of 35 distinct bile acids were quantified within murine ileal content (Figure 4C). When assessing the ileal bile acid profile, 3 bile acids, TUDCA, tauro-β-muricholic acid (TβMCA), and TCA were significantly different compared to pretreatment using a two-way ANOVA followed by Dunnett’s multiple comparisons post hoc test. For TUDCA, all three ursodiol treatments were significantly different from pretreatment (all treatments, p = 0.0001). For TβMCA, only the ursodiol 50 treatment was significantly different from pretreatment (p = 0.0001). For TCA, all three ursodiol treatments were significantly different from pretreatment (Ursodiol 50, p = 0.0002; Ursodiol 150, p = 0.0040, and Ursodiol 450, p = 0.0374). Within the ileal content, the two highest MDA scores from the Random Forest analysis were UDCA and TUDCA, with high concentrations of both these bile acids in the ursodiol 450 treatment group (Figure S2A). A Kruskal-Wallis one-way ANOVA test followed by Dunn’s multiple comparisons test was used to calculate the significance of an individual bile acid within each Ursodiol treatment group compared to pretreatment. For ileal content, UDCA, TUDCA, GUDCA, and LCA were significantly higher in ursodiol 450 treatment compared to pretreatment (p = 0.0007, p = 0.0013, p = 0.0022, and p = 0.0218, respectively; Figure S3A).

**Figure 4:**
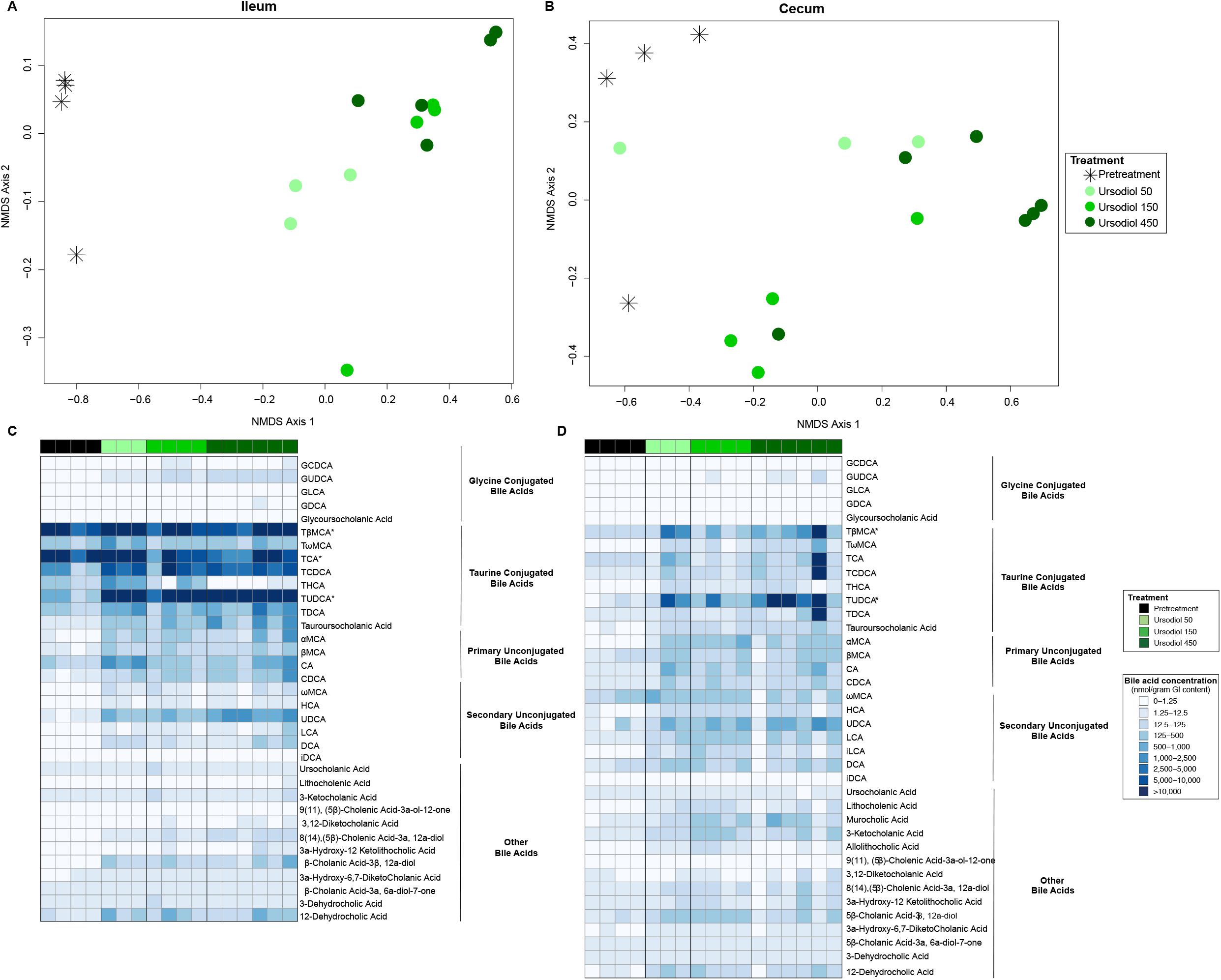
Alterations to the ileal and cecal bile acid metabolome associated with Ursodiol administration in conventional mice. NMDS ordination illustrates dissimilarity indices via Horn distances between bile acid profiles of paired **(A)** ileal and **(B)** cecal samples from pretreatment and Ursodiol treated mice. Statistical significance between Ursodiol treatment groups and pretreatment mice was determined by AMOVA. Targeted bile acid metabolomics of murine **(C)** ileal and **(D)** cecal content was performed by UPLC-MS/MS and identified 35 and 38 distinct bile acids respectively. Significance determined by a two-way ANOVA followed by Dunnett’s multiple comparisons post hoc test to compare comprehensive bile acid profiles of Ursodiol treatment groups to pretreatment (* denotes significance).

Cecal content bile acid profiles revealed segregation of the ursodiol treatments from pretreatment bile acid profiles (Figure 4B). A total of 38 distinct bile acids were quantified within murine cecal content (Figure 4D). When assessing the cecal bile acid profile, 2 bile acids, TUDCA and TβMCA were significantly different compared to pretreatment using a two-way ANOVA followed by Dunnett’s multiple comparisons post hoc test. For TUDCA, Ursodiol 50 and 450 treatment groups were significantly different from pretreatment (both treatments, p = 0.0001). For TβMCA, only the Ursodiol 50 treatment was significantly different from pretreatment (p = 0.0219). The two highest MDA scores from the Random Forest analysis were TCDCA and TUDCA, with high concentrations of both these bile acids in the Ursodiol 450 treatment group (Figure S2B). A Kruskal-Wallis one-way ANOVA test followed by Dunn’s multiple comparisons test was used to calculate the significance of an individual bile acid within each Ursodiol treatment group compared to pretreatment. For cecal content, LCA, 3-ketocholanic acid, and α-muricholic acid (αMCA) were significantly higher in the Ursodiol 150 treatment compared to pretreatment (p = 0.0143, p = 0.0255; and p = 0.0280, respectively; Figures S3B). UDCA, TUDCA, GUDCA, TβMCA, and MCA were significantly higher in the Ursodiol 450 treatment compared to pretreatment (p = 0.0.0307, p = 0.0047, p = 0.0160, p = 0.0352, and p = 0.0321, respectively; Figures S3B).

Serial fecal bile acid profiles revealed distinct segregation of the ursodiol treatments from each other and from pretreatment bile acid profiles (Figure 5A). A total of 38 distinct bile acids were quantified within murine feces (Figure 5B). When assessing fecal bile acid profiles, 4 bile acids, UDCA, TUDCA, MCA, and TβMCA were significantly different compared to pretreatment using a two-way ANOVA followed by Dunnett’s multiple comparisons post hoc test performed at each sampling day (Day 5, 8, 10, 12, and 15). Within the Ursodiol 50 treatment group, UDCA and TUDCA were significantly different from pretreatment only at Day 8 (p = 0.0296 and p = 0.0001, respectively). Within the Ursodiol 150 treatment group, UDCA and TUDCA were significantly different from pretreatment only at Day 15 (p = 0.0001 and p = 0.0107, respectively). Within the Ursodiol 450 treatment group, UDCA was significantly different from pretreatment at Days 5 (p =0.0020), 8 (p = 0.0007), 10 (p = 0.0044), and 15 (p = 0.0001). TUDCA was also significantly different from pretreatment in the Ursodiol 450 group at all sampling days (p = 0.0001 for all days). Additionally, MCA and TβMCA in the Ursodiol 450 treatment group on Day 15 were significantly different from pretreatment (p = 0.0001 for both).

**Figure 5:**
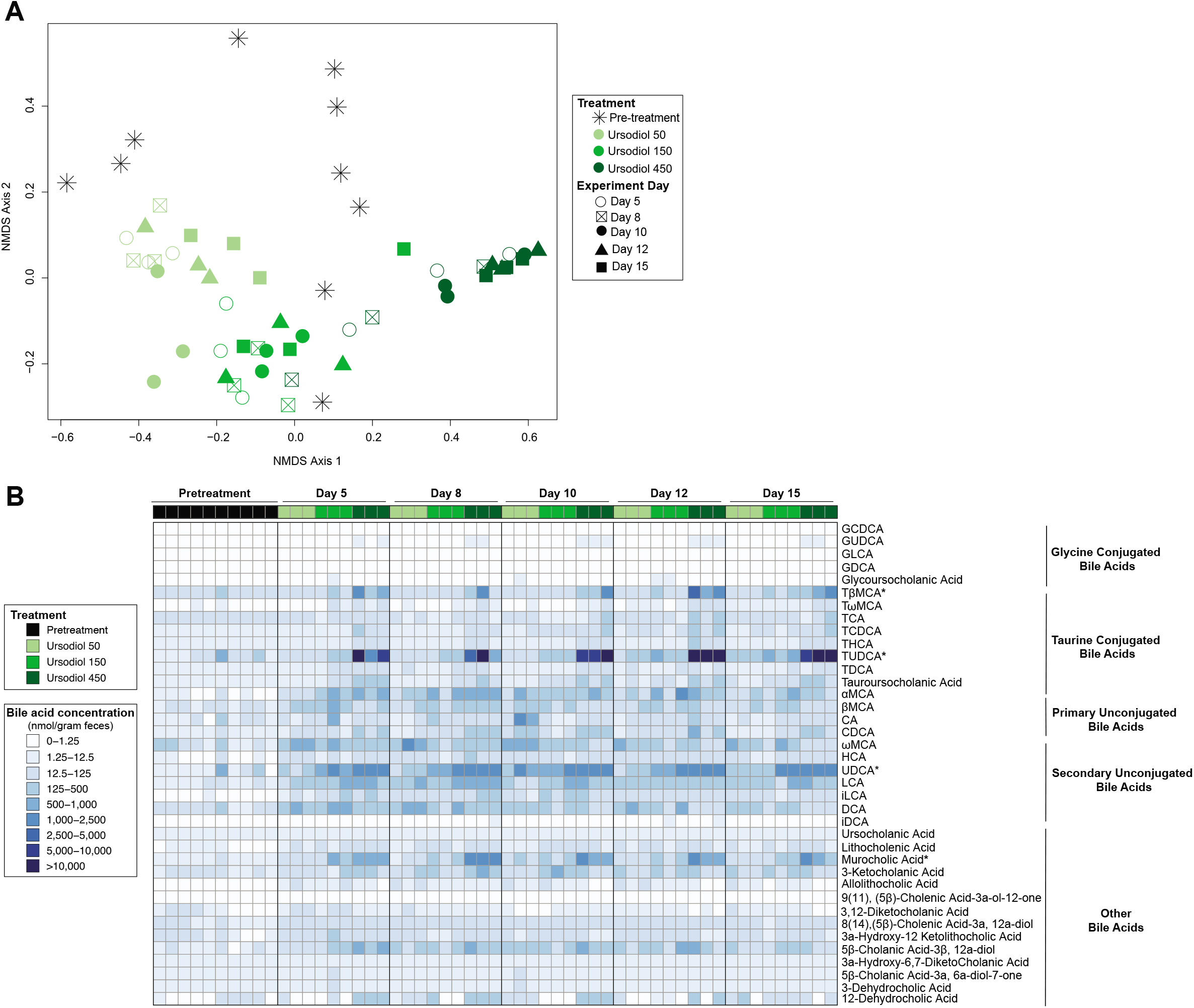
Alterations to the fecal bile acid metabolome throughout Ursodiol administration in conventional mice. **(A)** NMDS ordination illustrates dissimilarity indices via Horn distances between bile acid profiles of paired fecal samples. **(B)** Targeted bile acid metabolomics of murine feces was performed by UPLC-MS/MS and identified 38 distinct bile acids. Significance determined by a two-way ANOVA followed by Dunnett’s multiple comparisons post hoc test to compare comprehensive bile acid profiles of Ursodiol treatment groups to pretreatment (* denotes significance). Data represents two independent experiments (pretreatment, n = 10; n= 3 per treatment per sampling day).

Within serum, aside from a single ursodiol 50 treatment serum sample, the ursodiol treatments segregated distinctly from the pretreatment samples with Ursodiol treatments clustering together at day 21 (Figure S4A). A total of 35 distinct bile acids were quantified within murine serum samples (Figure S4B). The two highest MDA scores from the Random Forest analysis were TUDCA and UDCA, with high concentrations of both these bile acids in the Ursodiol 450 treatment group (Figure S4C). A Kruskal-Wallis one-way ANOVA test followed by Dunn’s multiple comparisons test was used to calculate the significance of an individual bile acid within each Ursodiol treatment group compared to pretreatment. UDCA, TUDCA, GUDCA, and LCA were significantly higher in Ursodiol 450 treatment compared to pretreatment (p = 0.0008, p = 0. 0007, p = 0.0230, and p = 0.0065, respectively; Figure S4D).

## Discussion

This study is the first to provide a comprehensive examination of how exogenously administered ursodiol shapes the gastrointestinal ecosystem in conventional mice. By evaluating the gut microbial community structure and bile acid pool throughout the gastrointestinal tract and in feces, we obtained a biogeographical view of ursodiol mediated ecological impact. Our findings indicate distinct ursodiol mediated alterations in the ileum, cecum, and feces likely attributed to biogeographical differences in the intestinal physiology and microbial ecology in each region.^39^

Dose dependent ursodiol mediated alterations in the gut microbial community structures were observed in the ileum and cecum (Figure 3). In both the ileum and cecum, members of the Lachnospiraceae family (phylum Firmicutes, Class Clostridia) significantly contributed to the observed alterations (Figure S1). Lachnospiraceae are Gram-positive obligate anaerobes, which are highly abundant in the digestive tracts of many mammals, including humans and mice.^40,41^ Members of the Lachnospiraceae have been linked to obesity^42–44^ and may provide protection from colon cancer,^45,46^ mainly due to their association with butyric acid production^47^, which is essential for microbial and host cell growth.^40^ Additionally, monocolonization of germfree mice with a Lachnospiraceae isolate resulted in greatly improved clinical outcomes and partial restoration of colonization resistance against the enteric pathogen *Clostridioides difficile*.^48^ Collectively, emphasizing the varied disease states where members of the Lachnospiraceae family are important and demonstrating potential applications of Ursodiol mediated Lachnospiraceae expansion to precisely modulate microbial mediated disease states.

Ursodiol administration resulted in global increases of several key bile acid species, namely UDCA, TUDCA, GUDCA, LCA, TCA, and TβMCA. Each of these bile acids can interact with bile acid activated receptors, including G-protein coupled bile acid receptor 5 (TGR5) and the farnesoid X receptor (FXR), and thus are able to regulate and alter host physiologic responses.^3–5^ Activation of either bile acid receptor has distinct physiologic consequences. For example, FXR regulates bile acid, glucose, and lipid homeostasis, and insulin signaling and immune responses.^3,4^ TGR5 regulates energy homeostasis, thermogenesis, insulin signaling, and inflammation.^3,4^ In terms of innate immune regulation, the overall response of FXR and TGR5 activation is maintenance of a tolerogenic phenotype within the intestine and liver (recently reviewed in Fiorucci *et al.*).^4^ Each bile acid species differ in their agonistic or antagonistic effects and affinity for FXR and TGR5 (see Table 1). This intensifies the complexity of unraveling the cumulative host physiologic responses resulting from ursodiol mediated bile acid metabolome alterations.

**Table 1:** Bile acid effects on bile acid receptors FXR and TGR5

| Bile Acid | Farnesoid X Receptor (FXR) | G-protein coupled bile acid receptor 5 (TGR5) |
| --- | --- | --- |
| UDCA | Antagonist | Agonist |
| TUDCA | Agonist<sup>54</sup> | Agonist<sup>53</sup> |
| GUDCA | Antagonist<sup>55</sup> | Agonist<sup>56</sup> |
| LCA | Agonist | Agonist |
| TCA | Agonist | Results in GLP-1 release<sup>57</sup> |
| TβMCA | Antagonist | - |

Table adapted from Wahlstrom et al., 2016<sup>3</sup> and Fiorucci et al., 2018<sup>4</sup> manuscripts

Additionally, bile acid species can directly and indirectly, through activation of the innate immune response, alter the gut microbial composition.^3,4^ Further adding to the interconnectedness and complexity of the gut microbiota-bile acid-host axis. Evaluation of the host intestinal transcriptome may elucidate local Ursodiol mediated impacts on host physiology and complete our examination of the gut microbiota-bile acid-host axis. Acquisition of such data, in combination with the comprehensive microbiome and bile acid metabolome data obtained in this study, could be integrated using bioinformatics and mathematical modeling to further illustrate these intricate interactions between the gut microbiota, bile acids, and the host in an Ursodiol altered intestinal ecosystem.

During ursodiol administration significant weight loss was noted in the ursodiol 50 and Ursodiol 450 treatments compared to untreated mice (Figure 2). We speculate that weight loss was attributed to bile acid TGR5 activation resulting in alteration to energy metabolism. A similar pathophysiology of weight loss attributed to bile acid activation of TGR5 is documented in patients following bariatric surgery.^49^ Circulating bile acids can activate TGR5 receptors within enteroendocrine cells, skeletal muscle, and brown adipose tissue.^50^ Aside from TGR5 mediated glucagon-like peptide-1 (GLP-1) release, which can improve glycemic control by increasing insulin secretion and sensitivity,^51^ TGR5 can facilitate weight loss by increasing resting energy expenditure by promoting conversion of inactive thyroxine (T4) into active thyroid hormone (T3).^52^ In our study, global large-scale increases in TUDCA, a TGR5 receptor agonist,^53^ were observed and may explain why weight loss occurred in our ursodiol treated mice. It is unclear why weight loss was not observed in the ursodiol 150 treatment group. Further investigation into TGR5 activation and subsequent modulation of energy expenditure with Ursodiol administration would be of interest.

In this study, we reported that daily ursodiol administration in conventional mice significantly impacts the gastrointestinal ecosystem, with alterations in the microbial composition and bile acid pool. Such substantial ecology changes are likely to modify host physiology. Ecological succession after ursodiol discontinuation was not evaluated in the present study, thus understanding how durable ursodiol mediated changes are in the mouse gastrointestinal systems remain unclear. Therefore, although ursodiol is generally well tolerated and safe to administer for various hepatic diseases,^6,19–27^ the long-term consequences of ursodiol mediated gastrointestinal ecologic shifts remains unknown. Further studies evaluating how exogenously administered bile acids, such as ursodiol, manipulate the dynamic gut microbiota-bile acid-host axis may elucidate how to restore health during disease states characterized by bile acid metabolism, including metabolic disease, obesity, IBD, and microbial-mediated colonization resistance against enteric pathogens such as *C. difficile.*

## Acknowledgements

JAW was funded by the Ruth L. Kirschstein National Research Service Award Research Training grant T32OD011130 by NIH. CMT is funded by the National Institute of General Medical Sciences of the National Institutes of Health under award number R35GM119438. This project was also funded by an intramural grant from the North Carolina State University College of Veterinary Medicine.

---

**Supplemental Figure 1: Lachnospiraceae family members significantly contribute to shifts in the microbial community seen with Ursodiol treatment in conventional mice. (A)** Ileal and **(B)** cecal principal coordinate analysis (PCoA) biplot using a Spearman correlation for top 10 significant OTUs. **(C)** Longitudinal fecal principal coordinate analysis (PCoA) biplot using a Spearman correlation for top 10 significant OTUs.

**Supplemental Figure 2: Bile acids that can differentiate between Ursodiol treatment groups. Variable-importance plot of the top 15 bile acids identified by Random Forest analysis in the (A)** ileum and **(B)** cecum. The mean accuracy value decrease (MDA score) is a measure of how much predictive power is lost if the given bile acid is removed or permuted in the Random Forest algorithm. Therefore, the more important a bile acid is to classifying samples into a treatment group, the further to the right the point is on the graph. Bile acid points are color-coded for relative concentrations of each bile acid within the Ursodiol 450 treatment group (red if their concentration is high in Ursodiol 450 treatment, gray if they were intermediate, and light blue if the concentrations were low). Each bile acid name is colored coded based on bile acid type (purple indicates glycine conjugated, orange indicates taurine conjugated, teal indicates primary unconjugated, blue indicates secondary unconjugated, and gray indicates other type of bile acid).

**Supplemental Figure 3: Alterations in the ileal and cecal bile acid metabolome associated with Ursodiol administration in conventional mice**. Box and whisker plots of **(A)** ileal and **(B)** cecal bile acids that were significantly altered in Ursodiol treated mice compared to pretreatment in any of the sample types evaluated (based on a Two-way ANOVA with Dunnett’s multiple comparisons post hoc test). Data represents two independent experiments (pretreatment, n = 4; Ursodiol 50, n = 3; Ursodiol 150, n = 4; Ursodiol 450, n= 6).

**Supplemental Figure 4: Alterations in the serum bile acid metabolome associated with Ursodiol administration in conventional mice. (A)** NMDS ordination illustrates dissimilarity indices via Horn distances between bile acid profiles of serum samples. **(B)** Targeted bile acid metabolomics of murine serum was performed by UPLC-MS/MS and identified 38 distinct bile acids. **(C)** Variable-importance plot of the top 15 bile acids identified by Random Forest analysis. Bile acid points are color-coded for relative concentrations of each bile acid within the Ursodiol 450 treatment group (red if their concentration is high in Ursodiol 450 treatment, gray if they were intermediate, and light blue if the concentrations were low). Each bile acid name is colored coded based on bile acid type (purple indicates glycine conjugated, orange indicates taurine conjugated, teal indicates primary unconjugated, blue indicates secondary unconjugated, and gray indicates other type of bile acid). **(D)** Box and whisker plots of bile acids that were significantly altered in Ursodiol treated mice compared to pretreatment in any of the sample types evaluated (based on a Two-way ANOVA with Dunnett’s multiple comparisons post hoc test). Data represents two independent experiments (pretreatment, n = 4; Ursodiol 50, n = 3; Ursodiol 150, n = 4; Ursodiol 450, n= 6).

