## Supplementary figures and images for "Secondary bile acid ursodeoxycholic acid (UDCA) alters weight, the gut microbiota, and the bile acid pool in conventional mice"

### Supplemental Figures

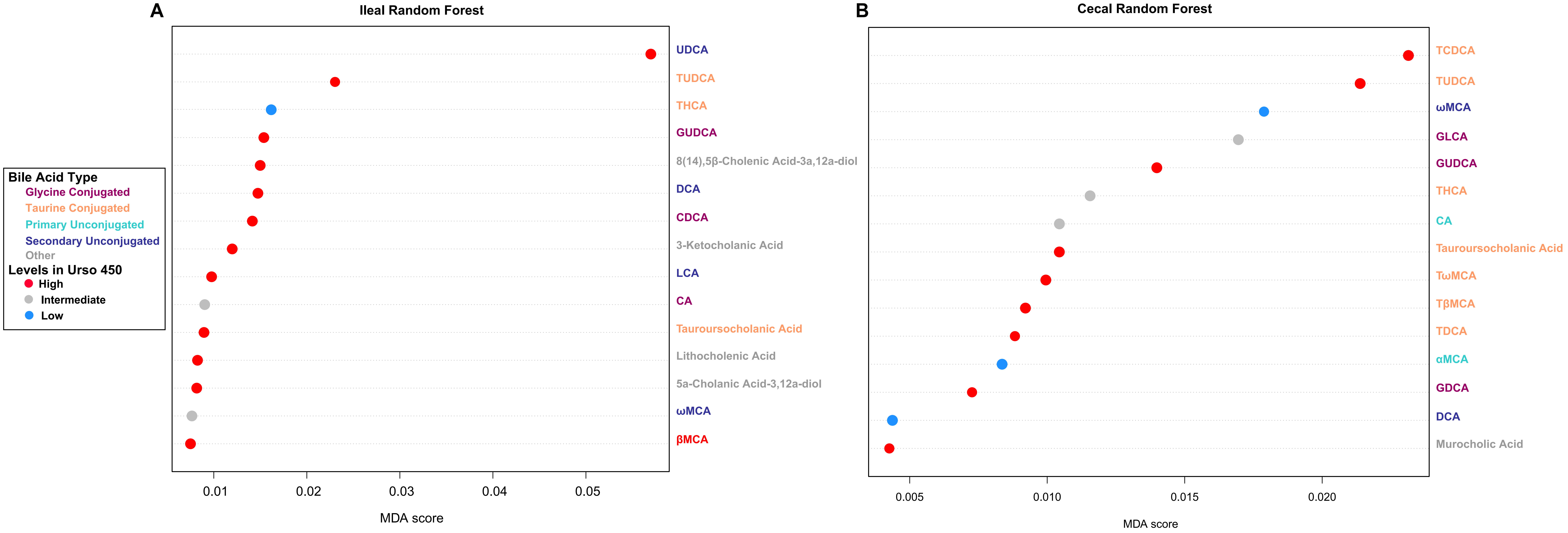

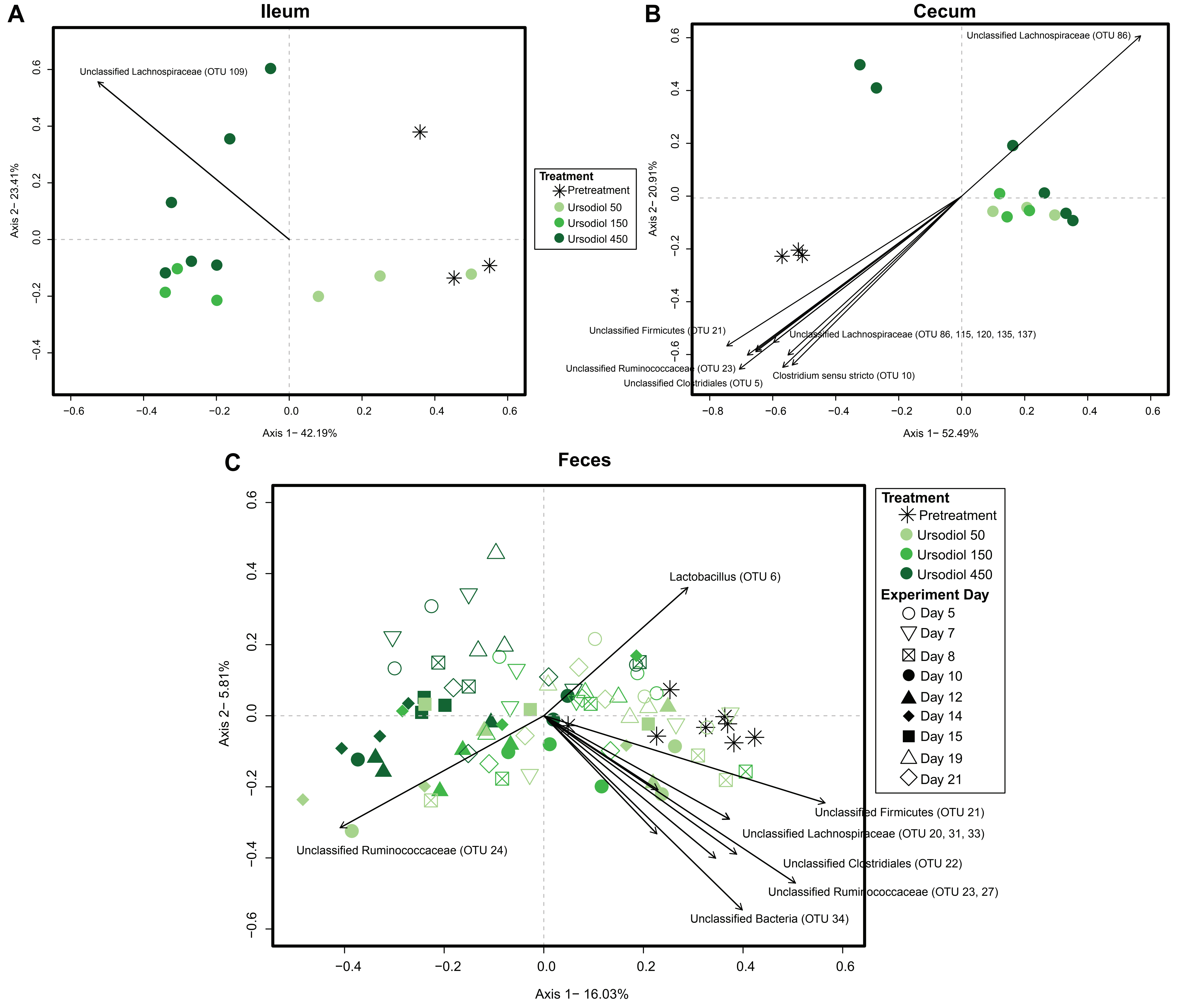

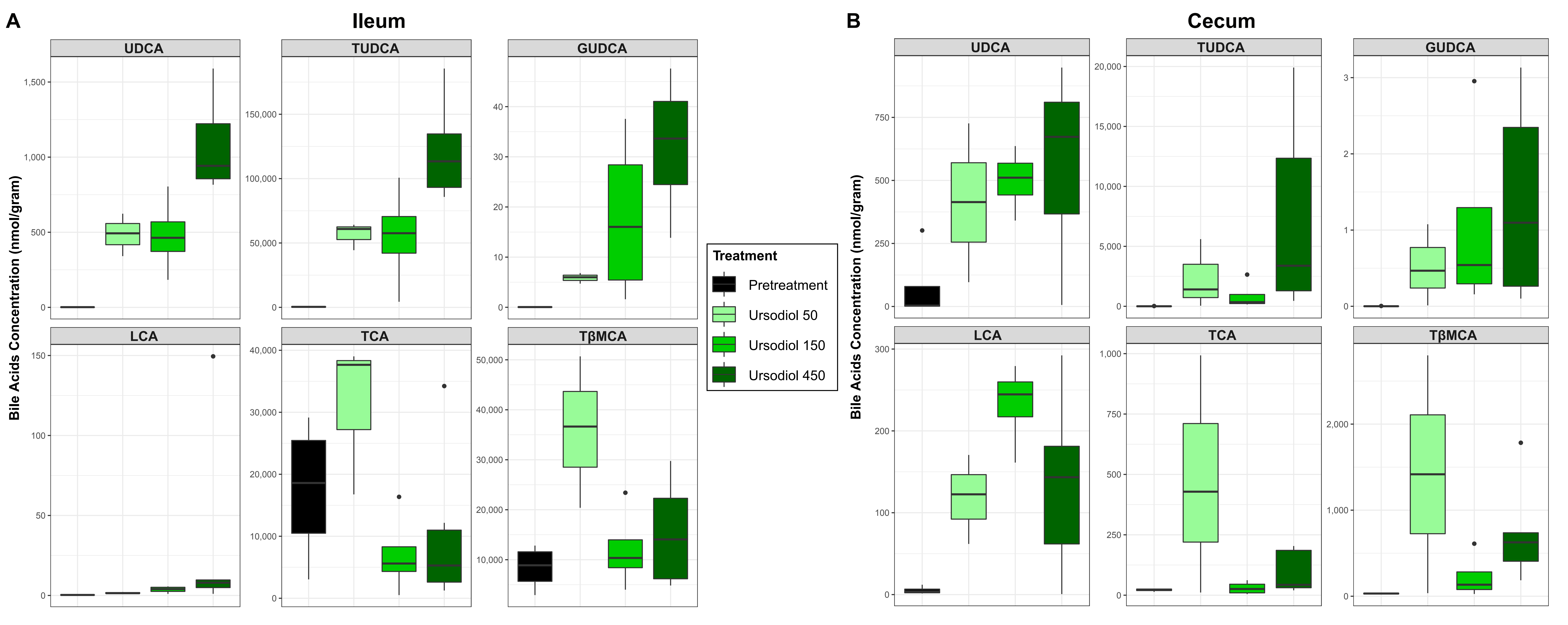

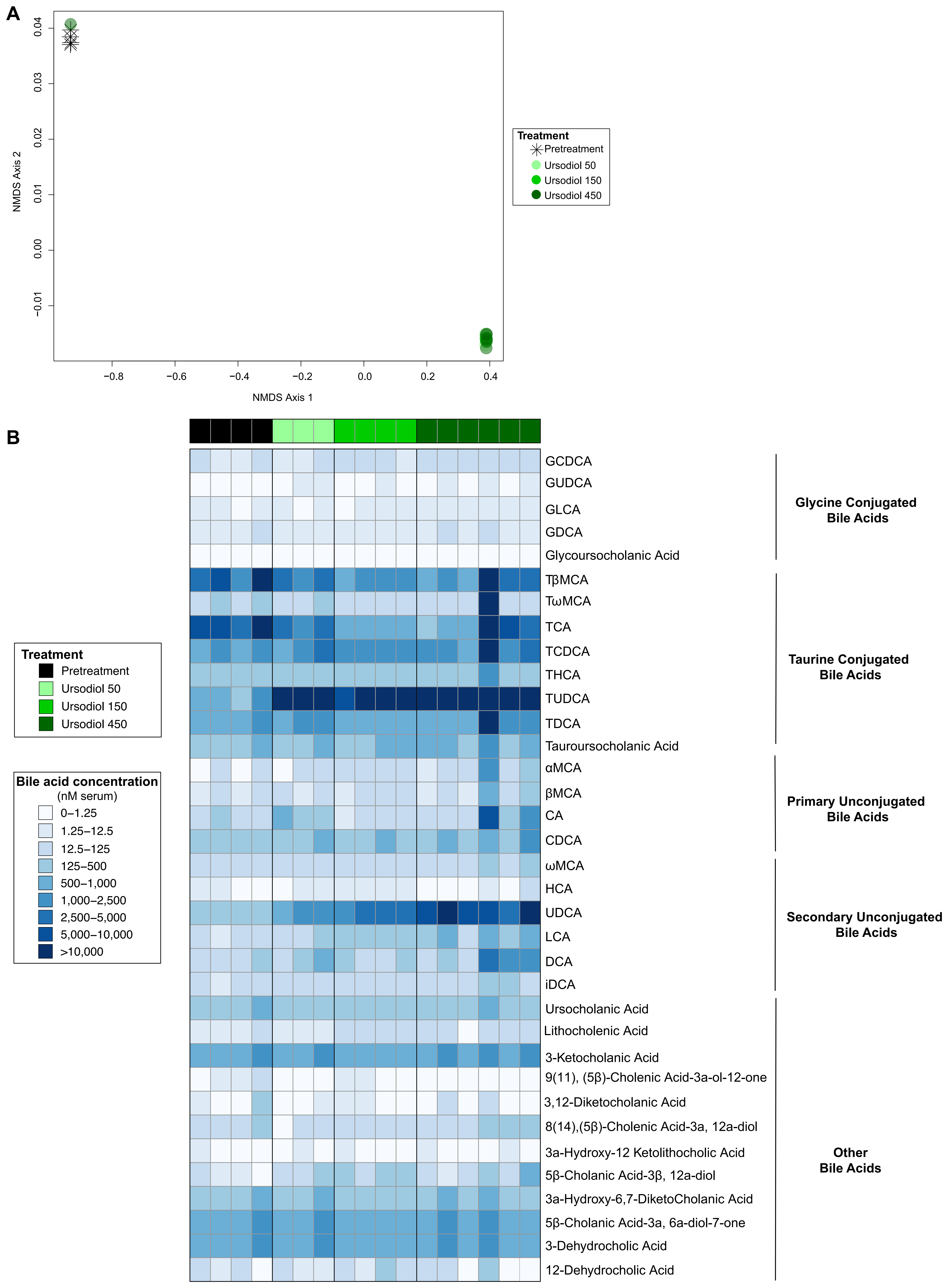

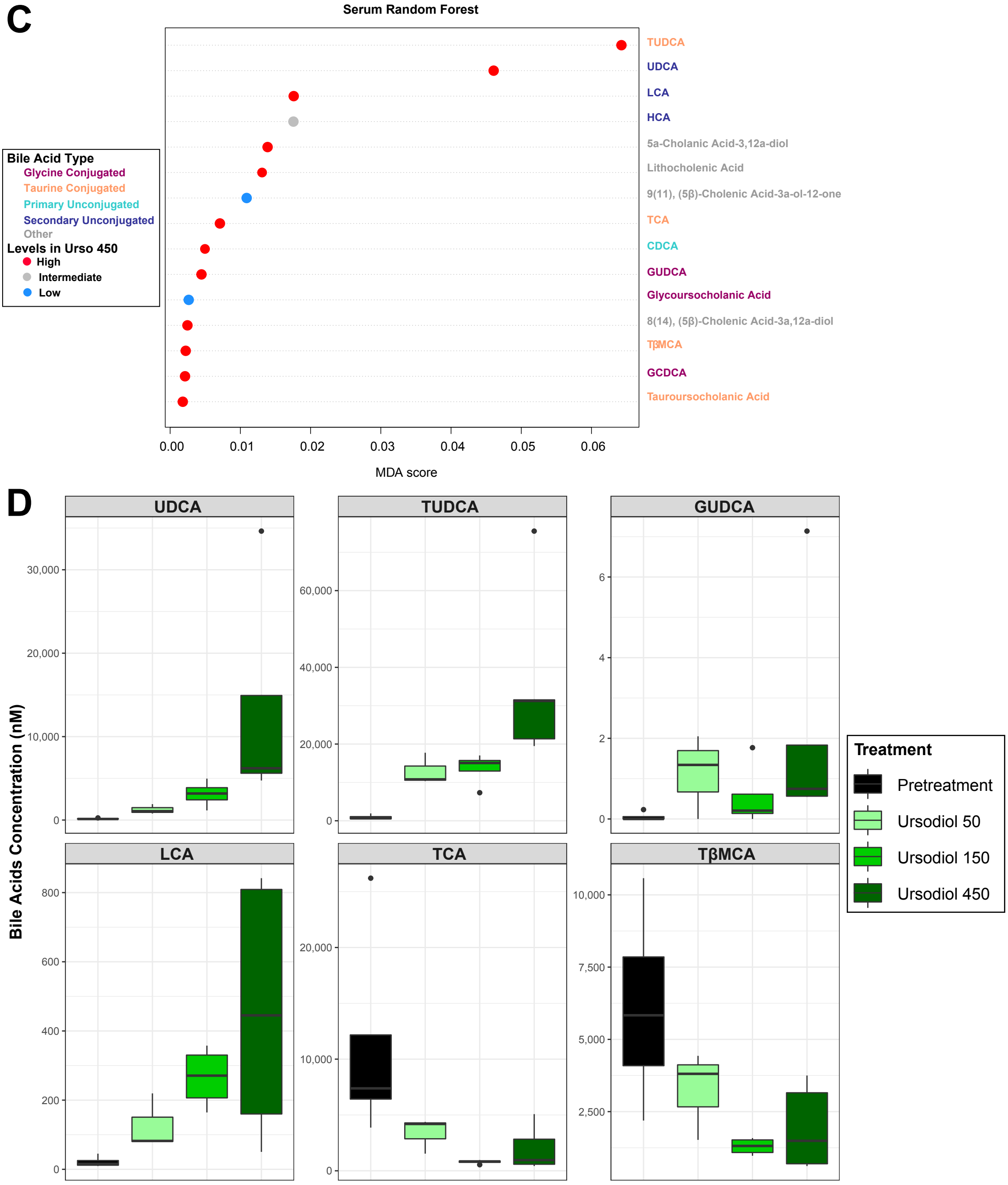
